# Co-targeting BCL-X_L_ and MCL-1 with DT2216 and AZD8055 synergistically inhibits small-cell lung cancer growth without causing on-target toxicities in mice

**DOI:** 10.1101/2022.09.12.507616

**Authors:** Sajid Khan, Patrick Kellish, Nick Connis, Dinesh Thummuri, Janet Wiegand, Peiyi Zhang, Xuan Zhang, Vivekananda Budamagunta, Nan Hua, Yang Yang, Umasankar De, Lingtao Jin, Weizhou Zhang, Guangrong Zheng, Robert Hromas, Christine Hann, Maria Zajac-Kaye, Frederic J. Kaye, Daohong Zhou

## Abstract

Small-cell lung cancer (SCLC) is an aggressive malignancy with limited therapeutic options. The dismal prognosis in SCLC is in part associated with an upregulation of BCL-2 family anti-apoptotic proteins, including BCL-X_L_ and MCL-1. Unfortunately, the currently available inhibitors of BCL-2 family anti-apoptotic proteins, except BCL-2 inhibitors, are not clinically relevant because of various on-target toxicities. We, therefore, aimed to develop an effective and safe strategy targeting these anti-apoptotic proteins with DT2216 (our platelet-sparing BCL-X_L_ degrader) and AZD8055 (an mTOR inhibitor) to avoid associated on-target toxicities while synergistically optimizing tumor response. Through BH3 mimetic screening, we identified a subset of SCLC cell lines that is co-dependent on BCL-X_L_ and MCL-1. After screening inhibitors of selected tumorigenic pathways, we found that AZD8055 selectively downregulates MCL-1 in SCLC cells and its combination with DT2216 synergistically killed BCL-X_L_/MCL-1 co-dependent SCLC cells, but not normal cells. Mechanistically, the combination caused BCL-X_L_ degradation and suppression of MCL1 expression, and thus disrupted MCL-1 interaction with BIM leading to an enhanced apoptotic induction. *In vivo*, DT2216+AZD8055 combination significantly inhibited the growth of cell line-derived and patient-derived xenografts and reduced tumor burden accompanied with extended survival in a genetically-engineered mouse (GEM) model of SCLC without causing significant thrombocytopenia or other normal tissue injury. Thus, these preclinical findings lay a strong foundation for future clinical studies to test DT2216+mTOR inhibitor combination in a subset of SCLC patients whose tumors are co-driven by BCL-X_L_ and MCL-1.

## INTRODUCTION

Small-cell lung cancer (SCLC) is a difficult-to-treat subtype of pulmonary carcinomas with a five-year survival rate of ~5% and median survival of less than a year^1,2^. First-line treatment for SCLC with combinations of a platinum-based agent (cisplatin or carboplatin) and etoposide (a topoisomerase-II inhibitor) has remained largely unchanged for almost 35 years. Recently, a triplet of atezolizumab (a PD-L1 antibody), carboplatin and etoposide, was approved as a first-line therapy for extensive-stage SCLC but provides only a moderate (~2 months) increase in overall survival compared to chemotherapy alone^3,4^. More importantly, the SCLC tumors are initially responsive to chemotherapy, however relapse occurs in majority (>80%) of cases, and there is no effective second-line treatment for relapsed SCLC^4,5^. There is thus an urgent and unmet need to find newer treatment strategies to effectively treat SCLC.

SCLC has historically been treated as a homogeneous disease. However, it has been recently demonstrated that both inter- and intra-tumoral heterogeneity occurs in SCLC, which is primarily responsible for the reduced therapeutic efficacy and resistance^6–9^. Since unique resistance mechanisms occur in response to specific therapies, specific oncoproteins and/or pathways need to be defined and targeted for effectively treating SCLC. Aberrant expression of anti-apoptotic BCL-2 family proteins has also been shown to account for intratumoral heterogeneity and therapeutic resistance in SCLC^6,10^. This has been therapeutically exploited with a dual BCL-X_L_/BCL-2 inhibitor ABT263 (navitoclax) that shows high efficacy in preclinical models of SCLC^11,12^. Unfortunately, the single agent efficacy of navitoclax against SCLC in phase-II clinical trials was limited^13^ because of the intratumoral heterogeneity and dependence of some of the cells within a tumor on other BCL-2 family proteins such as MCL-1. More importantly, the clinical use of ABT263 is hampered by dose-limiting severe thrombocytopenia caused by BCL-X_L_ inhibition^14,15^. We are now able to safely target BCL-X_L_ using proteolysis targeting chimeras (PROTACs, such as DT2216) without causing significant platelet toxicity (thrombocytopenia) as recently reported by our group^16–20^.

In this study, we aimed to develop a synergistic and safe therapeutic strategy for SCLC. This was achieved first by profiling the heterogeneity of survival dependence of a panel of SCLC cell lines on the BCL-2 family anti-apoptotic proteins by BH3 mimetic screening, where we used BH3 mimetics to target specific BCL-2 anti-apoptotic proteins followed by measurement of cell viability^21^. We defined a subset of SCLC cell lines that are co-dependent on BCL-X_L_ and MCL-1 for survival and focused on testing the ability to target these cell lines effectively and safely. Currently, such tumors cannot be safely targeted with available inhibitors. For example, MCL-1 inhibition causes severe cardiotoxicity and co-targeting BCL-X_L_ and MCL-1 with commercially available inhibitors further exacerbates normal tissue injury and causes lethality^22–24^. Therefore, we need to develop an alternate strategy to selectively suppress BCL-X_L_ and MCL-1 in co-dependent SCLC tumors to avoid their on-target and dose-limiting toxicities. We have devised a safer strategy to target these tumors as we found that BCL-X_L_/MCL-1 co-dependent SCLC cells can be synergistically killed with a combination of DT2216 (a selective BCL-X_L_ PROTAC degrader) and AZD8055 (an mTOR inhibitor). This was achieved by selective degradation of BCL-X_L_ and inhibition of MCL-1 expression in tumor cells with DT2216 and AZD8055, respectively, therefore avoiding on-target toxicity to platelets and other normal cells. *In vivo*, the combination of DT2216 and AZD8055 strongly inhibited growth of cell line-derived and patient-derived xenograft (PDX) tumors and reduced tumor burden as well as extended survival of a murine *Rb1/p53/p130* SCLC GEM model without causing significant thrombocytopenia or other normal tissue injury.

## MATERIALS AND METHODS

### Cell lines and culture

All the SCLC cell lines, except DMS53 and DMS114, were obtained from the original NCI-Navy Medical Oncology source supply^25^. DMS53 and DMS114 SCLC cell lines, CCD-18Co normal colon fibroblasts and WI-38 lung fibroblasts were purchased from the American Type Culture Collection (ATCC, Manassas, VA). SCLC cell lines were cultured in RPMI-1640 medium (Cat. No. 22400–089, Thermo Fisher, Waltham, MA). CCD-18Co and WI-38 cells were cultured in Dulbecco’s modified Eagle’s medium (DMEM) (Cat. No. 12430-062, Thermo Fisher). The culture media were supplemented with 10% heat-inactivated fetal bovine serum (FBS, Cat. No. S11150H, Atlanta Biologicals, GA), 100□U/mL penicillin and 100□μg/ml. streptomycin (Pen-Strep, Cat. No. 15140122, Thermo Fisher). Normal human bronchial epithelial (NHBE) cells were purchased from Lonza (Cat. No. CC-2541, Basel, Switzerland), and were cultured in Lonza’s bronchial epithelial cell growth medium (Cat. No. CC-3170) with supplements and growth factors (CC-4175). The stocks of NCI SCLC cell lines that we used were STR profiled by NIH, so the authenticity of the cell lines remained preserved. All cultures were confirmed for Mycoplasma negativity using the MycoAlert Mycoplasma Detection Kit (Cat. No. LT07–318). All the cell lines were maintained in a humidified incubator at 37 °C and 5% CO_2_.

### Chemical compounds

DT2216 was synthesized in Dr. Guangrong Zheng’s laboratory (University of Florida, Gainesville, FL) according to the previously described protocol^16^. AZD8055 (Cat. No. HY-10422) and everolimus (Cat. No. HY-10218) were purchased from MedChemExpress (Monmouth Junction, NJ). A1155463 (Cat. No. S7800), ABT199 (Cat. No. S8048), S63845 (Cat. No. S8383), and ABT263 (Cat. No. S1001) were purchased from SelleckChem (Houston, TX).

### Cell viability assays

The cell viability was measured by MTS assay according to the manufacturer’s protocol (Cat. No. G-111, Promega, Madison, WI) and as described previously^16,26^. EC_50_ values were determined using GraphPad Prism software (GraphPad Software, La Jolla, CA).

### BH3 mimetic screening

The screening was performed to determine the survival dependence of SCLC cells on different BCL-2 family anti-apoptotic proteins^21^. The cells were treated in 96-well plates with either a specific inhibitor of BCL-2 family anti-apoptotic proteins such as A1155463 (selective BCL-X_L_ inhibitor), venetoclax (selective BCL-2 inhibitor) and S63845 (selective MCL-1 inhibitor), or a non-specific inhibitor such as navitoclax (BCL-X_L_ BCL-2 dual inhibitor). Thereafter, the viability was assessed using MTS assay as described above.

### Co-immunoprecipitation

Cell pellets were lysed in the Pierce IP lysis buffer (Cat. No. 87787; Thermo Fisher) supplemented with protease and phosphatase inhibitors as described previously^16,26^. The supernatants were collected and precleared by incubating with 1 μg of mouse anti-IgG (Cat. No. sc-2025; Santa Cruz Biotechnology [SCB], Dallas, TX) and 20 μL of protein A/G-PLUS agarose beads (Cat. No. sc-2003; SCB) for 30 min at 4 °C. The supernatants containing 1 mg of protein were incubated with 2 μg of anti-MCL-1 (Cat. No. sc-12756; SCB) or anti-IgG antibody overnight followed by incubation with 25 μL protein A/G agarose beads for 1-2 h at 4 °C. Thereafter, the immunoprecipitates were collected by centrifugation, washed three times with IP lysis buffer, mixed with 50 μL of Laemmli’s SDS-buffer, denatured and then subjected to immunoblot analysis for BIM and MCL-1. Anti-rabbit HRP-conjugated Fc fragment specific secondary antibody (Cat. No. 111-035-046, dilution 1:10000, Jackson ImmunoResearch, West Grove, PA) was used to detect immune complexes in immunoblotting.

### Cell line-derived xenograft and PDX studies

All the animal procedures were performed in accordance with the rules of IACUC. CB-17 SCID-beige mice aged 5-6 weeks were purchased from the Charles River Laboratories (Wilmington, MA). NCI-H1048 (H1048) tumor cells at a density of 2.5×10^6^ per mouse in 50% Matrigel (Cat. No. 356237, Corning, Corning, NY) and PBS mixture were injected subcutaneously (s.c.) into the right flank region of the mice as described previously^16,26^. Tumor size was measured twice a week with digital calipers and tumor volume was calculated using the formula (Length×Width^2^×0.5). The mice were randomized into different treatment groups when the tumors reached ~150 mm^3^. Mice were treated with vehicle, AZD8055 (16 mg/kg, 5 days a week, p.o.), DT2216 (15 mg/kg, every four days [q4d], i.p.) and a combination of AZD8055 and DT2216. AZD8055 was formulated in 10% (v/v) DMSO, and 90% of 20% (w/v) Captisol (Cat. No. NC0604701, Cydex Pharmaceuticals a Ligand Company, San Diego, CA, USA) in normal saline and DT2216 was formulated in 50% phosal 50 PG, 45% miglyol 810N and 5% polysorbate 80. Mice were euthanized when they became moribund, or their tumor sizes reached a humane endpoint as per Institutional Animal Care and Use Committee (IACUC) policy. For euthanasia, animals were sacrificed by CO_2_ suffocation followed by cervical dislocation. The tumors were subsequently harvested, lysed and used for immunoblot analysis.

The PDX experiments were performed at the Johns Hopkins University. NOD-*scid* IL2Rgamma^null^ (NSG) mice aged 5-6 weeks were purchased from the Jackson Laboratory (Stock No. 005557, Bar Harbor, Maine, USA). LX47 SCLC PDX model was established and characterized by Dr. Christine Hann’s lab at the Johns Hopkins University and were propagated in NSG mice as reported previously^27^. The tumors were harvested after they reached 1500 mm^3^ and cut into ~2-3 mm fragments and were implanted s.c. after submerging in Matrigel in additional NSG mice^27^. The animals were randomized into different treatment groups when the tumors reached 100-200 mm^3^ and were treated as described above. The tumor volumes were normalized with their baseline readings on day-1 of treatment and are presented as normalized tumor volumes.

### Tumor induction and treatments in *Rb1/p53/p130* GEM model

The *Rb1^loxP/loxP^ p53^lox/loxP^ p130^lox2722/lox2722^* mouse strain was provided by Dr. Maria Zajac-Kaye (University of Florida)^28,29^. Presence of floxed sequences (loxP and lox2722) was confirmed by PCR as described previously^29^. Tumors were initiated when mice were 6–8 weeks of age by nasal inhalation of adenovirus serotype 5 expressing Cre-recombinase fused to enhancer GFP under a CMV promoter (Ad5CMVCre-eGFP, University of Iowa Vector Core, Cat. No. WC-U of Iowa 1174) at 2.5×10^7^ pfu/mouse. Drug treatments were started 110 days after Ad-Cre delivery, which was the timing to be required for the onset of SCLC tumorlets. Mice were treated with vehicle, AZD8055 (16 mg/kg, 5 days a week, p.o.), and a combination of DT2216 (15 mg/kg, q4d, i.p.) and AZD8055. Survival events were scored when the body condition score (as defined by the American Association for Laboratory Animal Science) of animals declined or per absolute survival events^30^. The lungs were excised upon euthanizing the individual mice, stained in Bouins’ solution and photographed as described previously^31,32^. The number of visible tumor nodules were counted with naked-eyes on the surface of each pair of lungs and are presented as average number of tumor nodules/lungs.

### Statistical Analysis

For analysis of the means of three or more groups, analysis of variance (ANOVA) test was performed. In the event that ANOVA justified post-hoc comparisons between group means, the comparisons were conducted using Tukey’s multiple-comparisons test. A two-sided unpaired Student’s *t*-test was used for comparisons between the means of two groups. *P* < 0.05 was considered to be statistically significant. The combination index (CI) was calculated using compusyn software (https://www.combosyn.com/). CI <1 indicates synergistic effect, CI = 1 indicates additive effect and CI > 1 indicates antagonistic effects. CDI <0.7 indicates significant synergistic effect.

## RESULTS

### BH3 mimetic screening identifies DT2216 sensitive and resistant SCLC cells *in vitro*

In order to establish the importance of the BCL-2 family of anti-apoptotic proteins in SCLC, we explored their mRNA expression in patient tumors via cBioPortal^33^. These data showed that *BCL2, BCL2L1* (BCL-X_L_ coding gene) and *MCL1* are highly expressed in most SCLC patient tumors. Interestingly, the BCL-X_L_ mRNA levels were consistent in all the tumor samples (**Fig. 1a**). Similarly, immunoblot analysis also suggests that these proteins are abundantly expressed in most SCLC cell lines (**Supplementary Fig. 1**). Next, we systematically evaluated the effects of inhibitors of BCL-2 family anti-apoptotic proteins, i.e., A1155463 (a selective BCL-X_L_ inhibitor), venetoclax (a selective BCL-2 inhibitor), S63845 (a selective MCL1 inhibitor) and navitoclax (a BCL-X_L_/BCL-2 inhibitor) on the viability of a panel of 20 SCLC cell lines, which was named as the BH3 mimetic screening^21^. The cell lines with EC_50_ <1 μM for a particular inhibitor were considered sensitive. SCLC cell lines showed differential dependence on different BCL-2 family anti-apoptotic proteins, with a majority of them depending primarily on BCL-X_L_ (8/20 or 40%), while some depended on a combination of BCL-X_L_ and BCL-2 (3/20 or 15%), and only a few depended primarily on BCL-2 (1/20 or 5%) or MCL-1 (1/20 or 5%), as indicated by their sensitivity to these specific inhibitors (**Fig. 1b**). One cell line (H211) was found to be sensitive to all of the tested inhibitors indicating that all three, i.e., BCL-X_L_, BCL-2 and MCL-1 are equally important for its survival. These findings are corroborated with the transcriptomic profiling suggesting that BCL-X_L_ is one of the highly expressed BCL-2 family anti-apoptotic protein, and therefore, a promising therapeutic target in SCLC tumors^10–12^. Further, we evaluated the activity of our BCL-X_L_ PROTAC degrader DT2216 on the viability of these SCLC cell lines where 50% (10/20) cell lines were found to be sensitive to DT2216 with an EC_50_ of <1 μM. All of the cell lines that were found to be sensitive to BCL-X_L_ inhibitor were sensitive to DT2216 as well. Interestingly, H1059 cell line was not sensitive to BCL-X_L_ inhibitor but was found to be sensitive to DT2216. Interestingly, six of the cell lines did not respond to any of these inhibitors or DT2216 suggesting that these may co-depend on a combination of either BCL-X_L_/MCL-1 or BCL-2/MCL-1 or BCL-X_L_/BCL-2/MCL-1 (**Fig. 1b**). Overall, these results suggest that BCL-X_L_ plays a crucial role in SCLC cell survival and DT2216 is a broad-spectrum antitumor agent than a BCL-X_L_ inhibitor in killing BCL-X_L_-dependent SCLC cells.

**Figure 1.**
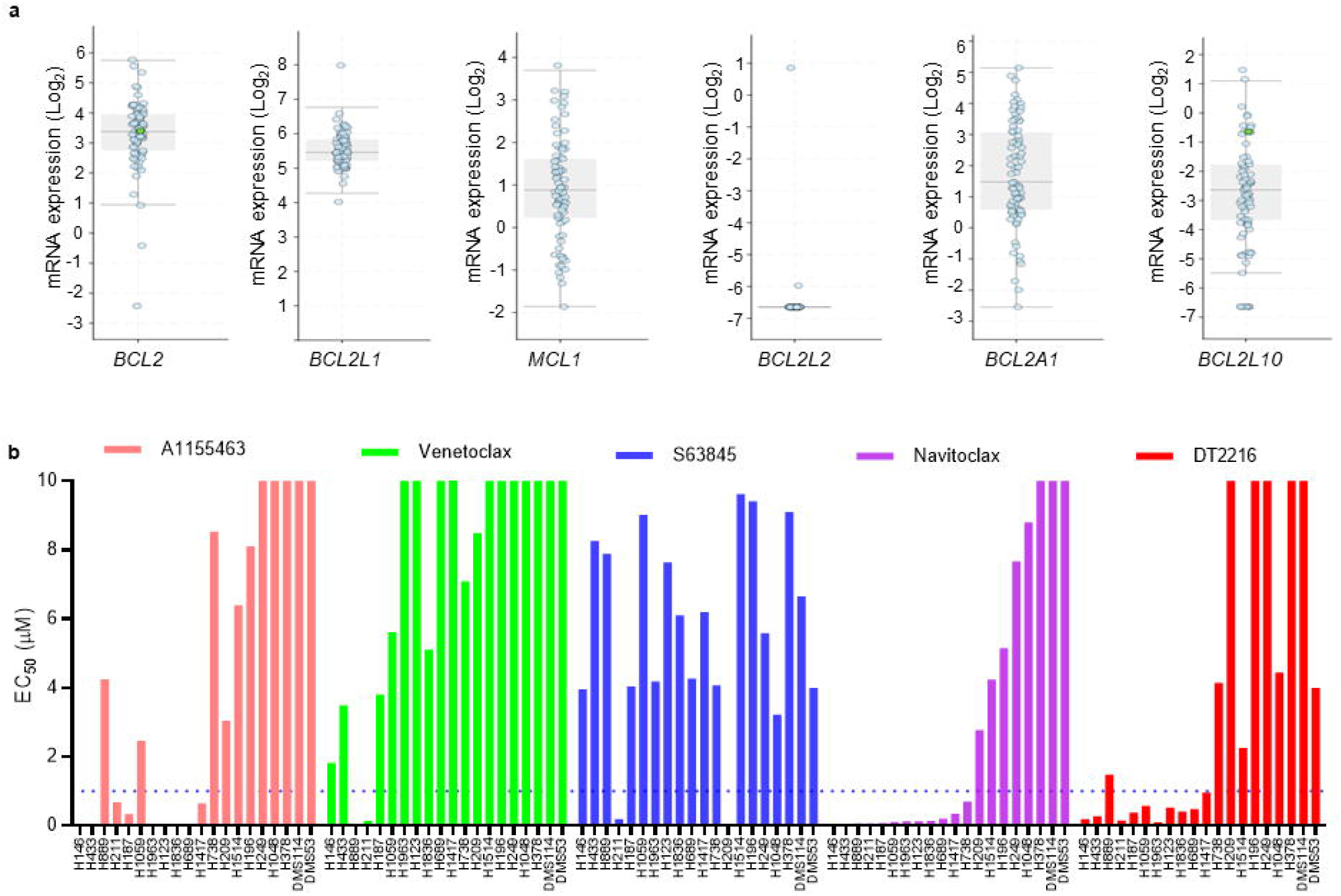
The SCLC cell lines show differential survival dependence on BCL-2 family proteins. **a,** The mRNA expression of BCL2 family anti-apoptotic genes as obtained from U Cologne study via cBioPortal^31^. **b,** The EC_50_ values of A1155463 (a selective BCL-X_L_ inhibitor), venetoclax (a selective BCL-2 inhibitor), S63845 (a selective MCL-1 inhibitor), navitoclax (a BCL-X_L_/BCL-2 dual inhibitor) and DT2216 (a selective BCL-X_L_ PROTAC degrader) were derived from % viability of indicated human SCLC cell lines after they were incubated with increasing concentrations of these inhibitors for 72 h. When an inhibitor showed an EC_50_ of ≤1 μM in any given cell line, the cell line was considered as sensitive to that inhibitor and dependent on its target BCL-2 family protein.

### A combination of MCL-1 inhibitor with BCL-X_L_ inhibitor or PROTAC is not selective to tumor cells

In the BH3 mimetic screening, SCLC cell lines showed varying responses to the inhibitors of different BCL-2 family anti-apoptotic proteins. Based on these dependencies, we divided SCLC cell lines into different categories; 1) primarily dependent on BCL-X_L_ (such as H1963), 2) mainly dependent on BCL-2 (such as H889), 3) mostly dependent on MCL-1 (such as H209), 4) BCL-X_L_ and BCL-2 co-dependent (such as H1059), 5) BCL-X_L_ and MCL-1 co-dependent (such as H378), and 6) BCL-X_L_, BCL-2, and MCL-1 dependent (such as H211) (**Fig. 2a**). When two or more proteins were needed to be simultaneously inhibited in order to sensitize a particular cell line, we called it co-dependent on those proteins. For example, H1059 cells were only sensitive upon simultaneous inhibition of BCL-X_L_ and BCL-2 using navitoclax, therefore, we categorized them as BCL-X_L_ and BCL-2 co-dependent. These results demonstrate that SCLC cells are heterogeneous in their dependencies on BCL-2 anti-apoptotic proteins.

**Figure 2.**
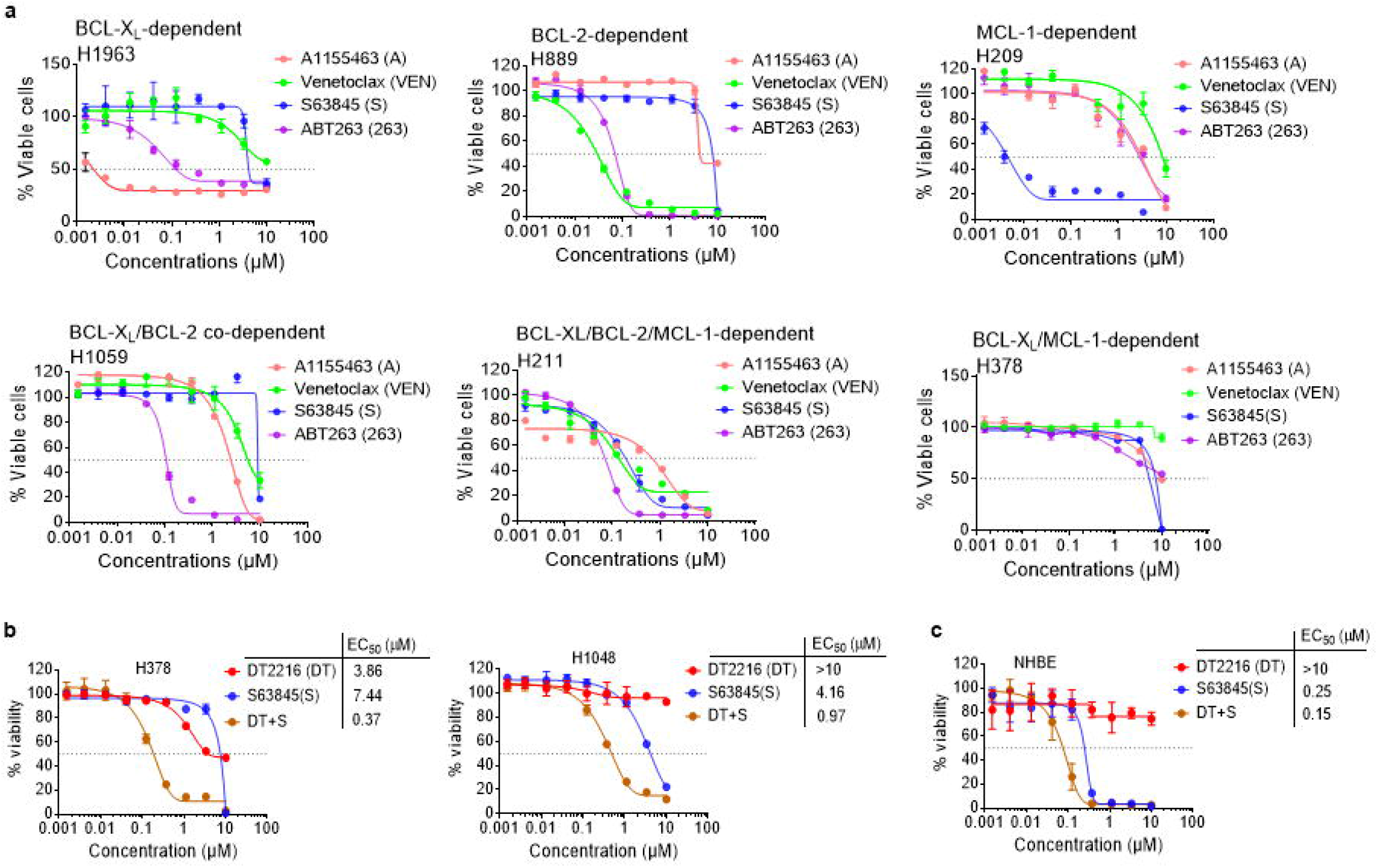
A subset of SCLC cell lines is co-dependent on BCL-X_L_ and MCL-1. **a,** Viability graphs of representative SCLC cell lines showing dependence on individual or a combination of BCL-2 family anti-apoptotic proteins. The data are presented as mean ± SD (*n* = 3 replicate cell cultures). **b, c,** Viability of H378 and H1048 SCLC (b) and NHBE (c) cells after they were treated with increasing concentrations of DT2216 (DT) or S63845 (S) or their combination (1:1 ratio) for 72 h. EC_50_ values for individual agents and combinations are shown.

Next, we focused on targeting BCL-X_L_ and MCL-1 co-dependent cells, as these type of SCLC tumors are highly resistant to chemotherapy and cannot be safely eradicated with the currently available inhibitors of BCL-2 family proteins because the combination of BCL-X_L_ and MCL-1 inhibitors causes severe normal tissue toxicities and lethality in mice^23,24,34^. This was confirmed by our study showing that a combination of DT2216 and MCL-1 inhibitor (S63845) synergistically kills not only tumor cells (**Fig. 2b**), but also normal cells from different tissue origins such as human bronchial epithelial cells (NHBE), colon fibroblasts (CCD-18Co) and lung fibroblasts (WI38) (**Fig. 2c; supplementary Fig. 2a**). These findings rule out the possibility to use the combination of a BCL-X_L_ inhibitor/PROTAC and an MCL-1 inhibitor to treat BCL-X_L_ and MCL-1 co-dependent SCLC in the clinic.

### DT2216+AZD8055 combination synergistically kills BCL-X_L_/MCL-1 co-dependent SCLC cells through degradation of BCL-X_L_ and suppression of MCL-1 expression, respectively, in a tumor cell-selective manner

Next, we sought to identify a strategy to selectively suppress MCL-1 expression in SCLC cells by rationally targeting certain tumorigenic proteins/pathways. We screened several clinical-stage compounds and FDA-approved drugs including doxorubicin (a topoisomerase II inhibitor and a commonly used chemotherapeutic), SNS-032 (a promiscuous cyclin-dependent kinase [CDK] inhibitor that primarily targets CDK9), AZD8055 (an ATP-competitive catalytic mTORC1/2 inhibitor), TD-19 (a cancerous inhibitor of protein phosphatase 2A [CIP2A] inhibitor) and piperazine (PPZ, a protein phosphatase 2A [PP2A] activator). All of these therapeutic agents have previously been shown to suppress MCL-1 expression through different mechanisms ^10,34-37^. Here, we aimed to determine whether any of these agents could selectively suppress MCL-1 expression in tumor cells while having minimal/no significant effect on the expression of MCL-1 in normal cells. In agreement with previous reports, doxorubicin, SNS-032 and AZD8055 caused significant suppression of MCL-1 expression in all tested SCLC cell lines (**Fig. 3a; Supplementary Fig. 2b**). Among these inhibitors tested, only AZD8055 showed minimal effect on the expression of MCL-1 in normal cells (**Fig. 3b; Supplementary Fig. 2c**). None of these agents significantly altered BCL-X_L_ or BCL-2 levels in both tumor as well as normal cells. Further, we analyzed the dose-dependent effect of AZD8055 on the expression of MCL-1, BCL-X_L_ and BCL-2 in H378 and H1048 SCLC and NHBE normal cells. AZD8055 caused dose-dependent suppression of MCL-1 expression in tumor cells without any effect in normal cells. As expected, AZD8055 had no observable effects on the expression of BCL-X_L_ and BCL-2. Also, AZD8055 dose-dependently inhibited activation of the mTOR downstream targets, i.e., p-4EBP1 and p-S6, indicating an on-target effect (**Fig. 3c, d**).

**Figure 3.**
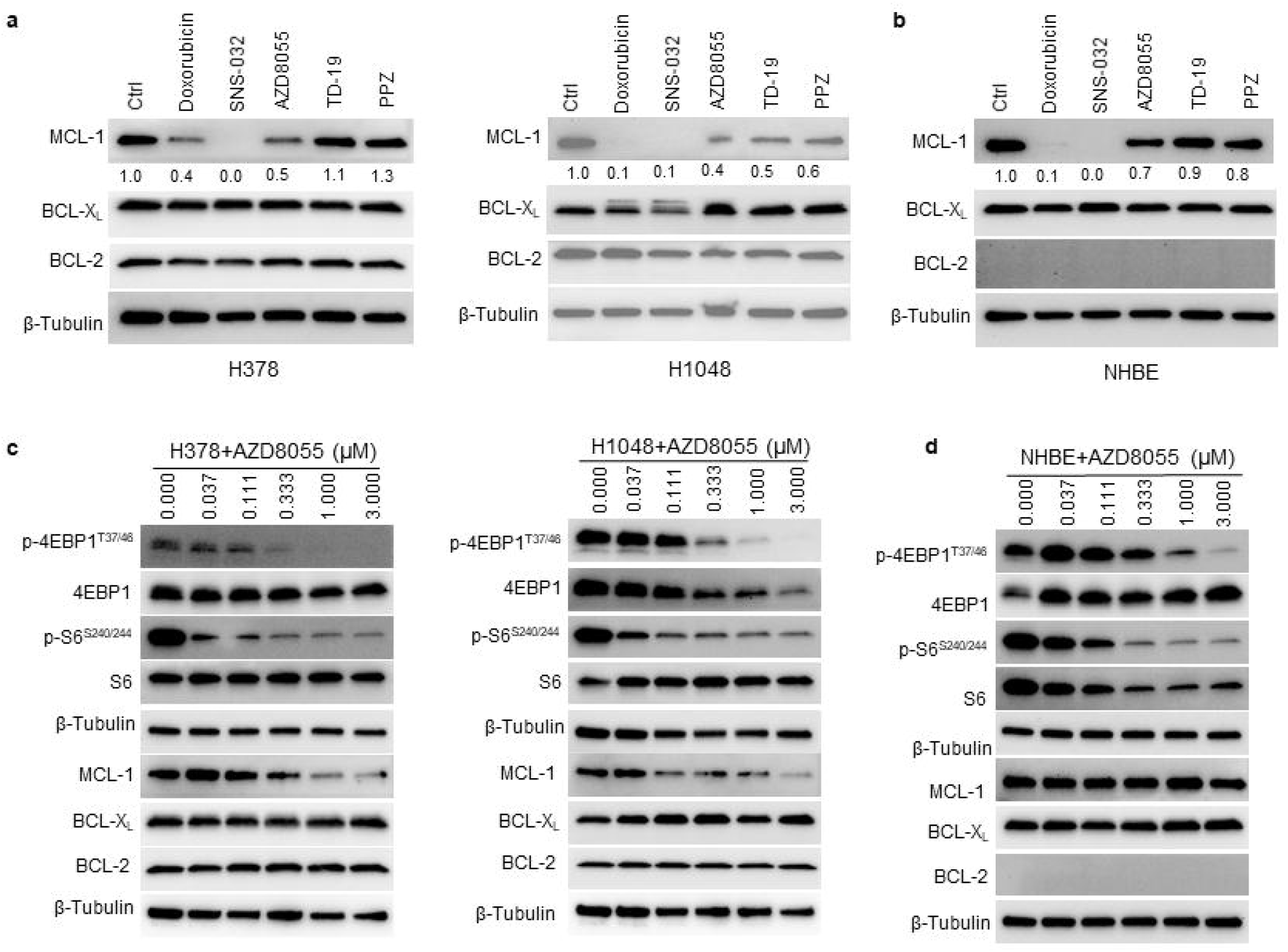
mTOR inhibitor AZD8055 selectively suppresses MCL-1 expression in tumor cells. **a, b,** Immunoblot analyses of MCL-1, BCL-X_L_ and BCL-2 in SCLC H378 and H1048 cell lines (a) and NHBE cells (b) after they were treated with indicated agents (1 μM each) for 24 h. Normalized densitometric values for MCL-1 blots are shown underneath. **c, d,** Immunoblot analysis of mTOR substrates (p-4EBP1 and p-S6), MCL-1, BCL-X_L_ and BCL-2 after 24 h treatment with indicated concentrations of AZD8055 in H378 and H1048 SCLC cell lines (c) and NHBE cells (d).

Since AZD8055 was found to selectively suppress MCL-1 expression in SCLC cells, we next evaluated it in combination with DT2216 on the viability of BCL-X_L_/MCL-1 co-dependent SCLC cells as well as normal cells. The combination was found to synergistically inhibit the viability of H378 and H1048 SCLC cells, but not normal NHBE and CCD-18Co cells (**Fig. 4a, b; Supplementary Fig. 2d**). DT2216 treatment induced a dose-dependent BCL-X_L_ degradation in H1048 cells as expected. Because 1 μM of DT2216 was found to completely deplete BCL-X_L_, we selected this concentration for further *in vitro* experiments (**Supplementary Fig. 3**). We found that the combination of DT2216 and AZD8055 synergistically inhibit the colony formation and induce apoptosis in H1048 cells (**Supplementary Fig. 4a, b**). Furthermore, DT2216+AZD8055 combination profoundly induced PARP cleavage compared to individual agents in tumor cells, but not in NHBE cells, further confirming tumor cell-selectivity of the combination (**Fig. 4c, d**).

**Figure 4.**
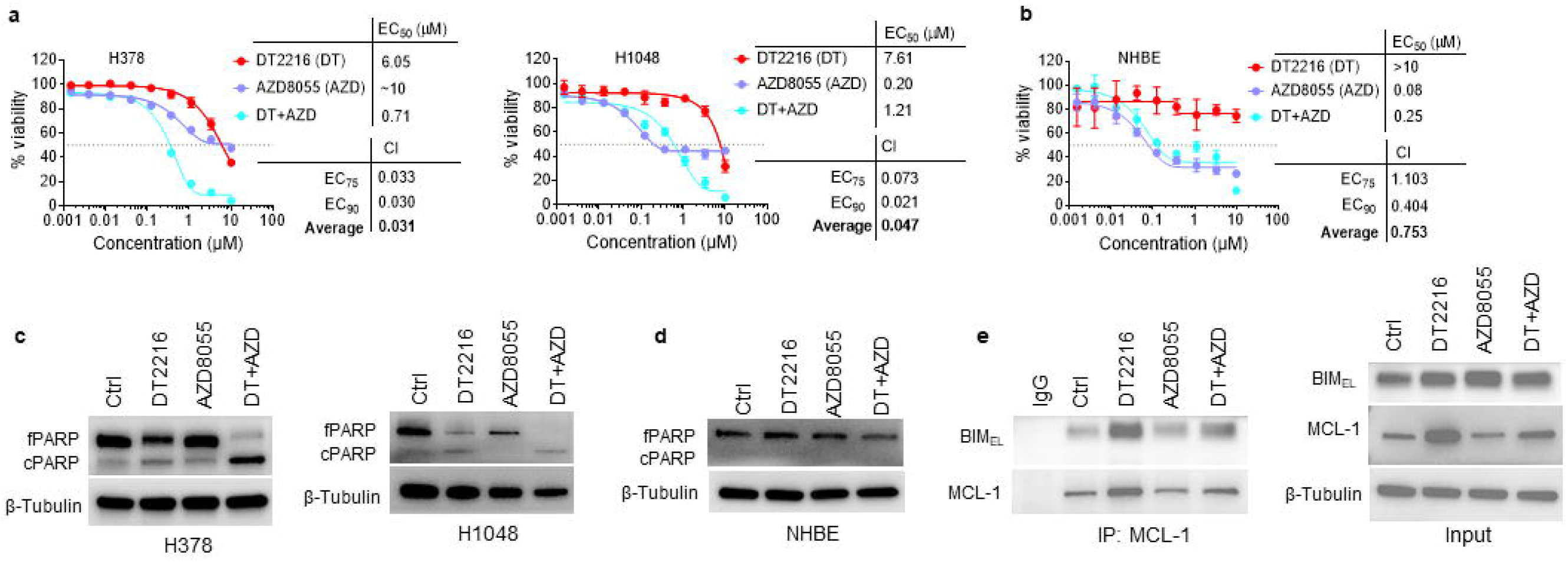
DT2216+AZD8055 combination synergistically kills BCL-X_L_/MCL-1 co-dependent SCLC cells through disruption of MCL-1/BIM interaction. **a, b,** Viability of H378 and H1048 SCLC (a) and NHBE (b) cells after they were treated with increasing concentrations of DT2216 (DT) or AZD8055 (AZD) or their combination (1:1 ratio) for 6 days. EC_50_ and CI values are shown. The data presented in **a** and **b** are mean ± SD (*n* = 3 replicate cell cultures). **c, d,** Immunoblot analyses of full-length PARP (fPARP) and cleaved PARP (cPARP) in SCLC H378, H1048 (c) and NHBE (d) cell lines after they were treated with 1 μM each of DT2216 or AZD8055 or their combination for 24 h. **e**, Immunoprecipitation analysis of MCL-1 followed by immunoblotting of BIM and MCL-1 in H1048 cells after they were treated with 1 μM each of DT2216 and/or AZD8055 for 24 h.

Unlike AZD8055 that inhibits both mTORC1 and mTORC2, everolimus is a rapamycin derivative and a selective mTORC1 inhibitor. Everolimus is FDA-approved for use as an immunosuppressant and for treating certain tumors. We wondered whether everolimus can also synergistically kill BCL-X_L_/MCL-1 co-dependent SCLC cells when combined with DT2216. Indeed, the combination of everolimus and DT2216 synergistically reduced viability and induced apoptosis in H378 and H1048 cells (**Supplementary Fig. 5a, b)**. This was accompanied with a significant reduction in MCL-1 expression by everolimus (**Supplementary Fig. 5c**). These results suggest that the inhibition of mTORC1 may be primarily responsible for the suppression of MCL-1 expression.

To gain further insights into the mechanism of synergistic activity of DT2216+AZD8055, we performed co-immunoprecipitation (co-IP) analysis of MCL-1 and BIM because BIM acts as a BH3 only pro-apoptotic protein that can trigger apoptosis by activating the apoptotic effectors BAX and BAK and the sequestration of BIM by MCL-1 or BCL-X_L_ inhibits apoptosis^38^. Since BIM is known to bind both BCL-X_L_ and MCL-1, we hypothesized that the degradation of BCL-X_L_ with DT2216 may lead to increased association of BIM with MCL-1, and co-treatment with DT2216+AZD8055 may disrupt this interaction. In line with our hypothesis, we found that DT2216 treatment increases the BIM/MCL-1 association which was disrupted when the cells were treated with DT2216+AZD8055 combination (**Fig. 4e**).

### DT2216+AZD8055 combination synergistically inhibits tumor growth in SCLC xenograft and PDX models

To further assess the combined mTOR inhibition and BCL-X_L_ degradation as a potential therapeutic strategy for BCL-X_L_ and MCL-1 co-dependent SCLC, we first investigated the efficacy of DT2216+AZD8055 combination using the H1048 xenograft tumor model. In agreement with the *in vitro* results, DT2216 alone had no effect on H1048 tumor growth. Although AZD8055 alone shows some efficacy, the effect was not significant. On the other hand, the combination of DT2216 and AZD8055 led to a significant inhibition of tumor growth (**Fig. 5a**). More importantly, the combination treatment appeared to be safe as indicated by no significant change in mouse body weights after the treatment (**Fig. 5b**). In addition, we did not observe any gross organ toxicities upon necropsy in mice treated with DT2216+AZD8055. Furthermore, the combination caused no clinically significant decrease in platelets (**Fig. 5c**). Next, we confirmed that DT2216 and AZD8055 led to a significant reduction in BCL-X_L_ and MCL-1 levels, respectively, which was associated with a significant inhibition of mTOR substrates (p-4EBP1 and p-S6) upon AZD8055 treatment in H1048 tumor lysates (**Fig. 5d, e**).

**Figure 5.**
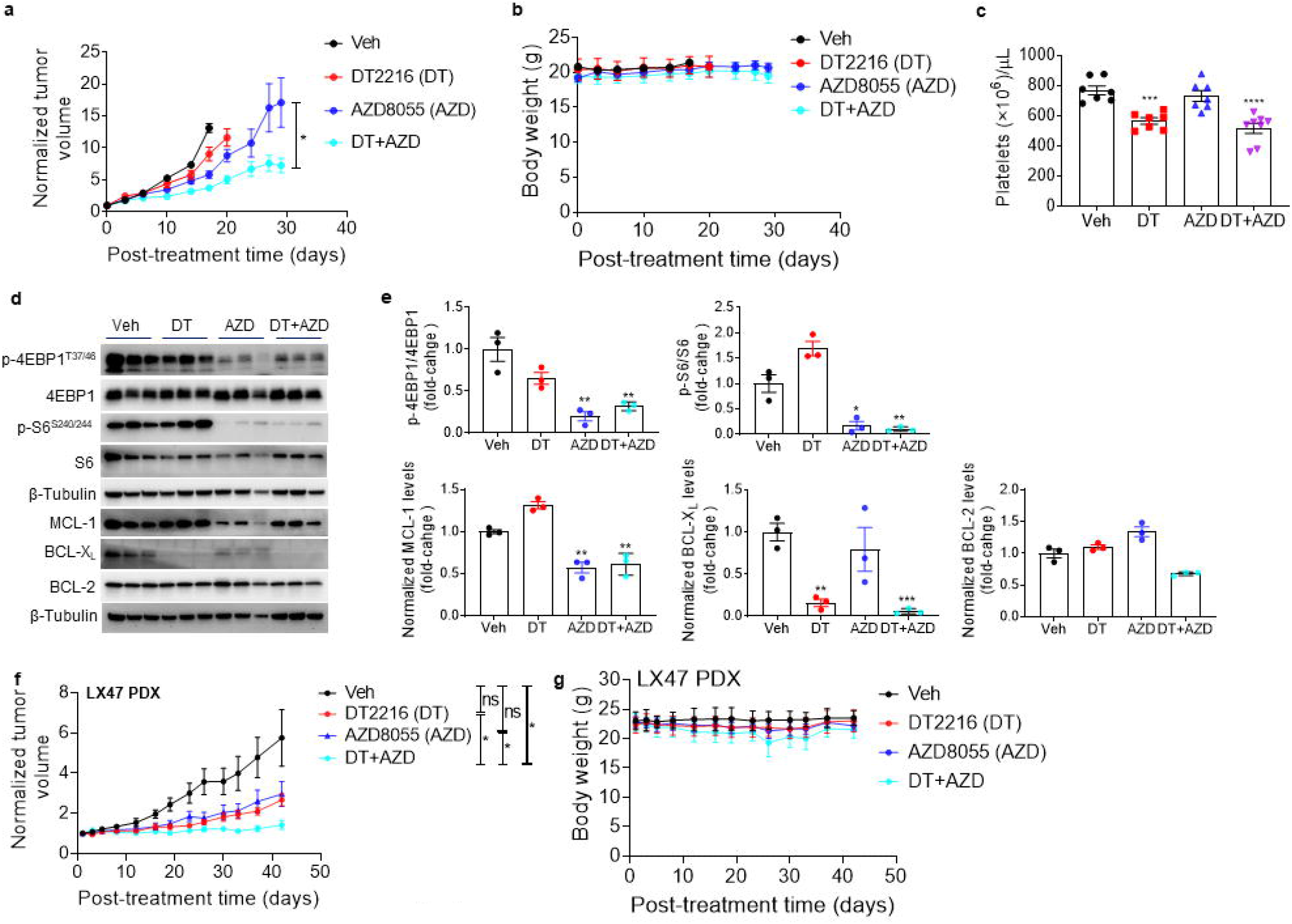
The combination of DT2216 and AZD8055 has stronger antitumor responses in H1048 xenograft and SCLC PDX models. **a, b,** Tumor volume (a) and mouse body weight (b) changes in H1048 xenografts after treatment with vehicle (veh), DT2216 (DT, 15 mg/kg/q4d, i.p.) or AZD8055 (AZD, 16 mg/kg/5x week, p.o.) or a combination of DT and AZD. Data are presented as mean ± SEM (*n* = 5-6 mice at the start of treatment). **c**, Platelet counts 24 h after first dose of DT2216 (DT) and/or AZD8055 (AZD) in H1048 xenograft mice as in **a**. **d,** Immunoblot analysis of indicated proteins in H1048 xenograft tumors at the end of treatments as in **a** (*n* = 3 mice per group). **e**, Densitometry of immunoblots shown in panel **d**. **f, g**, Tumor volume (f) and mouse body weight (g) changes in LX47 PDX tumors after treatment with veh, DT2216 (DT, 15 mg/kg/q4d, i.p.) or AZD8055 (AZD, 16 mg/kg/5x week, p.o.) or a combination of DT and AZD. Data are presented as mean ± SEM (*n* = 8-9 mice at the start of treatment).

Since the PDX models better recapitulate human cancers *in vivo*, we next evaluated the effect of the combination of DT2216+AZD8055 in the LX47-SCLC PDX model. We found that the combination significantly inhibited the tumor growth in this PDX model of SCLC, whereas single agents failed to have significant effects. Moreover, the tumor growth inhibition with the combination was significant as compared to single agent DT2216 or AZD8055 treatment (**Fig. 5f**). Again, we did not observe any significant body weight loss with the combination treatment in these mice (**Fig. 5g**).

### DT2216+AZD8055 combination inhibits lung tumor growth and extends mice survival in the *Rb1/p53/p130* GEM model

Next, we used the conditional-mutant *Rb1/p53/p130* mouse model of SCLC to test the efficacy of DT2216+AZD8055 combination. In these mice,*Rb1, Tp53* and *p130* tumor suppressor genes, have flanking lox sequences which are under control of Cre recombinase. These genes are deleted upon administration of Adenovirus (Ad5CMVCre-eGFP), specifically in the lung epithelial cells and lead to tumor formation in the lungs^28,29^. Since the *Rb1/p53/p130* mouse tumors closely resemble human SCLC, it is a suitable preclinical mouse model to evaluate newer therapeutics against SCLC. We determined a mean tumor onset time of 110 days after administration of adenovirus titer of 2.5×10^7^ pfu/mouse to induce the formation of small visible tumor nodules in the mouse lungs. Therefore, we started treating the mice after 110 days with DT2216 and/or AZD8055 for up to 150 days (**Fig. 6a**). We did not include DT2216 group because DT2216 was not found to have any significant effect on tumor growth when used alone in this model in a previous study as shown in Fig. 6e. Mice were euthanized, and the survival was recorded at the humane endpoint when the body condition score of animals declined^30^. At the end of the experiment i.e, 150 days after treatment, 2/8 (25%), 2/8 (25%) and 4/7 (~57%) mice were alive in vehicle, AZD8055 and combination-treatment groups, respectively. The median survival time was 131, 149.5 and >166 days in vehicle, AZD8055 and the combination-treatment group, respectively (**Fig. 6b**). We excised the lungs from these mice and observed that the combination-treated lungs had much smaller tumor nodules as compared to vehicle or AZD8055 treatments (**Fig. 6c**). Moreover, the number of total tumor nodules per lungs was also significantly reduced in the combination-treated mice (**Fig. 6d**). The histopathological staining in the lungs further evidenced the reduction in tumor burden in the combination-treated mice (**Fig. 6e**). We did not see any significant changes in the lungs, spleen, liver, or kidney weights after combination treatment (**Supplementary Fig. 6a**), neither did we see any clinically significant reduction in platelets and other blood cell counts (**Supplementary Fig. 6b, c**). Overall, these results suggest that the DT2216+AZ8055 combination is a promising therapeutic strategy against BCL-X_L_/MCL-1 co-dependent SCLC, and therefore this combination could potentially be tested in clinical trials in future.

**Figure 6.**
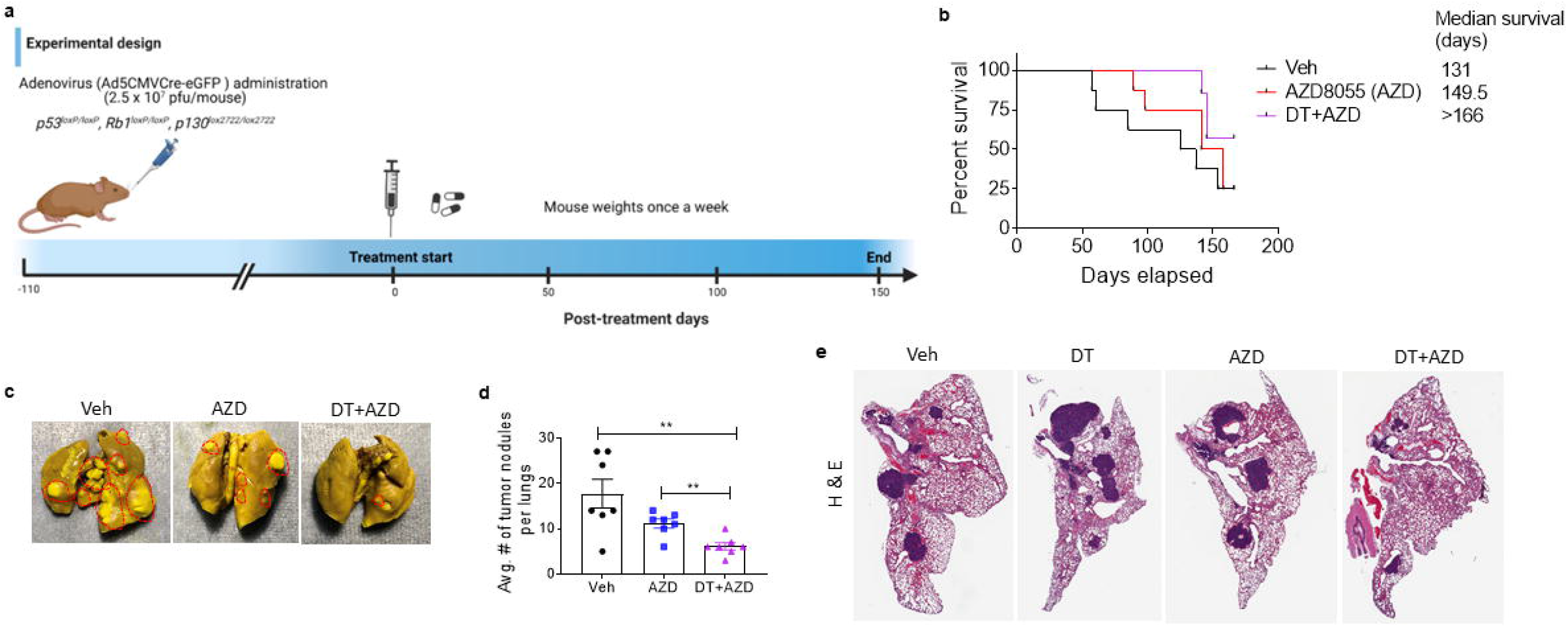
The combination of DT2216 and AZD8055 has stronger antitumor responses as compared to individual agents in a GEM model of SCLC. **a,** Experimental design of *Rb1/p53/p130* GEM model. Mice were administered with 2.5×10^7^ pfu/mouse of Ad5CMVCre-eGFP through nasal inhalation. Treatment was started 110 days post-infection and continued until day 150. Specifically, the mice were treated with vehicle (veh), AZD8055 (AZD, 16 mg/kg/5x week, p.o.) and AZD (16 mg/kg/5× week, p.o.) + DT2216 (DT, 15 mg/kg/q4d, i.p.) (*n* = 8, 8 and 7 mice in veh, AZD and combo groups, respectively) **b,** Kaplan-Meier survival analysis of mice as treated in **a**. The median survival time is shown on the right. At the end of experiment (day 166), 2, 2 and 4 mice were alive in veh, AZD and DT+AZD groups, respectively. **c**, Bouin’s-stained represented images of excised lungs from veh, AZD and DT+AZD groups. The tumor nodules in the lungs are circled. As shown, the tumor nodules were much smaller in the lungs of combination-treated mice as compared to veh or AZD-treated mice. **d**, Number of tumor nodules were counted in both dorsal and ventral sides of the lungs and the average number of tumor nodules per mouse lungs in each group are shown (mean ± SEM, *n* = 7-8). Statistical significance was determined by Student’s *t*-test. \*\**p* <0.01. **e**, Hematoxylin and eosin (H & E) staining of representative lung sections in veh, DT, AZD and DT+AZD groups from a separate experiment in which the mice were treated with veh, DT (15 mg/kg/q4d, i.p.), AZD (16 mg/kg/5× week, p.o.), and a combination of DT and AZD for 50 days. The mice were euthanized 24 h after last treatment.

## DISCUSSION

SCLC shows high inter- and intra-tumoral heterogeneity which is primarily responsible for the reduced therapeutic efficacy and resistance^6–9^. This heterogeneity is in part attributed to an aberrant expression of anti-apoptotic BCL-2 family proteins which are crucial for the survival of SCLC cells^10–12^. In the present study, we first established survival dependence of SCLC cell lines on different BCL-2 family anti-apoptotic proteins using BH3 mimetic screening. We found that most SCLC cell lines are dependent on BCL-X_L_ for their survival and are highly sensitive to a BCL-X_L_ inhibitor or our recently developed BCL-X_L_ PROTAC DT2216 (now a clinical candidate in phase-I trial)^16,17^. On the other hand, some SCLC cell lines were found to be dependent on multiple co-expressed BCL-2 family proteins such as concurrent BCL-X_L_ and BCL-2, or concurrent BCL-X_L_ and MCL-1. BCL-X_L_/BCL-2 co-dependent cell lines could be targeted with navitoclax, however, the clinical translation of navitoclax is hampered by dose-limiting on-target toxicity of thrombocytopenia^14,15^. More importantly, BCL-X_L_/MCL-1 co-dependent tumor cells cannot be safely targeted with the currently available inhibitors because the combination of BCL-X_L_ and MCL-1 inhibitors causes severe tissue damage and lethality in mice^22–24^. Therefore, finding a strategy to selectively target BCL-X_L_ and MCL-1 in tumors without causing significant on-target toxicity was the primary goal of our current study for treating an important difficult-to-treat subset of SCLC.

Since systemic MCL-1 inhibition with currently available inhibitors leads to normal tissue toxicity and lethality when combined with a BCL-X_L_ inhibitor/PROTAC, we sought to identify a strategy to selectively suppress MCL-1 expression in SCLC tumor cells. After screening several compounds that target oncogenic pathways known to upregulate MCL-1^10,34–37^, we identified that AZD8055, an mTORC1/2 ATP-competitive inhibitor and everolimus, a selective mTOR1 inhibitor, can selectively suppress MCL-1 expression in SCLC cells but not in normal cells. In contrast, other compounds such as doxorubicin and SNS-032 non-specifically downregulated MCL-1 in tumor as well as in normal cells. This was correlated with high basal mTOR activation in MCL-1 dependent SCLC cell lines (data not shown). The mTOR activation has been shown to enhance cap-dependent MCL-1 translation, and therefore, inhibition of mTOR suppresses MCL-1 at post-translational level ^34,36,37,39^. Of note, the combination of DT2216 with AZD8055 or everolimus synergistically inhibited the viability and induced apoptosis in BCL-X_L_ and MCL-1 co-dependent SCLC cell lines, but not the normal cells from different tissue origins including lungs and colon.

Our further investigation using different *in vivo* SCLC models including conventional xenografts, PDX and *Rb1/p53/p130* GEM model strongly suggests that the combination of DT2216 and AZD8055 can effectively inhibit the SCLC growth in mice. More importantly, the combination was found to be safer as evidenced by no significant decrease in mouse body weights as well as no clinically significant reduction in different blood cells including platelets, and the absence of any observable tissue pathology. Interestingly, results using the SCLC GEM model, that closely recapitulates genetic heterogeneity found in human SCLC, indicate that the combination of DT2216+AZD8055 strongly inhibits the tumor growth in the lungs and extended the survival of mice. The effectiveness of the combination in GEM model was encouraging because the tumors in this model have been shown to be highly resistant to cisplatin plus etoposide doublet chemotherapy^34^.

These findings have important clinical implications because tumor-specific targeting of BCL-X_L_ and MCL-1 has high therapeutic value. Two prior studies have found that a combination of navitoclax and AZD8055 synergistically kills different tumor cells from SCLC and *BRAF* or *KRAS*-mutated colorectal cancer^34,36^. Moreover, a study from Dr. Christine Hann’s group has demonstrated synergy between rapamycin and ABT737 (a predecessor compound of navitoclax) against different PDX models of SCLC^27^. In another study, the combined inhibition of BCL-X_L_ and mTOR was found to synergistically induce apoptosis in *PIK3CA*-mutated breast cancer^39^. However, these studies did not comprehensively evaluate tumor-selectivity and on-target toxicities of this combination (especially in mouse models) and given that BCL-X_L_ inhibitors including navitoclax cause thrombocytopenia, a combination of navitoclax with AZD8055 (or another mTOR inhibitor) may not be clinically feasible^14,15^. In contrast, by combining AZD8055 with platelet-sparing BCL-X_L_ degrader DT2216, we achieved tumor-selective targeting of both MCL-1 and BCL-X_L_, respectively, without significant on-target toxicity both in cell culture and in mice. This approach can also be applied to other tumor types which are co-dependent on MCL-1 and BCL-X_L_^26,40^. One limitation of our *in vitro* BH3 mimetic screening is that it requires live cells, so it cannot be used to stratify patients who will benefit from the combination. In that case dynamic BH3 profiling assay would be more useful as demonstrated previously^41,42^.

More recently, SCLC tumors have been classified into four molecular subtypes based on the relative expression of four transcriptional regulators, i.e., ASCL1, NEUROD1, POU2F3, and YAP1^5,7^. Each of these molecular subtypes show distinct therapeutic vulnerabilities^8,9^. For example, ASCL1-high subtype (classic or neuroendocrine (NE) SCLC)^7^ has been shown to be sensitive to BCL-2 inhibition with ABT263^43^. Interestingly, we observed that most variant-SCLC cells (non-NE consisting of POU2F3 and/or YAP1 subtype)^7^ were dependent on co-expression of BCL-X_L_ and MCL-1. Although, these subtypes are not being used to direct treatment decisions for SCLC at present, we predict that POU2F3 and/or YAP1 SCLC subtypes could be better targeted with a combination of DT2216 and an mTOR inhibitor. In this case, immunohistochemical or proteomics profiling of four of these transcriptional regulators could be a better alternative to BH3 profiling for stratifying patients who can specifically benefit from DT2216+mTOR inhibitor combination. In the current study we used AZD8055 as a proof-of-concept, but it would be suitable to use an FDA-approved mTOR inhibitor (such as everolimus) for combination with DT2216 in clinical trials.

In conclusion, the findings from our *in vitro* and *in vivo* studies suggest that the combination of DT2216+AZD8055 is tumor-selective and synergistic against BCL-X_L_/MCL-1 co-dependent subset of SCLC and is tolerable in mice without significant on-target toxicity and normal tissue injury. These findings have high potential for near-term clinical translation as the combination of DT2216 and an FDA-approved mTOR inhibitor (such as everolimus) can be rapidly evaluated in SCLC patients, given that DT2216 is already in phase-I clinical trial (Identifier: NCT04886622). This approach can help to effectively treat an important and difficult-to-treat subset of SCLC patients in the near future.

## Acknowledgements

Funding from US National Institutes of Health (NIH) grant R01 CA242003 (D.Z. & G.Z.), R01 CA241191 (D.Z. & G.Z.), R01 CA219836 (D.Z.), R01 CA260239 (D.Z., W.Z. & G.Z.) and Florida Department of Health grant 8JK04 (F.J.K. and M.Z-K).

## Author contributions

S.K. conceived, designed, and supervised the study, performed most of the *in vitro* and *in vivo* experiments, analyzed and interpreted data, and wrote and revised the manuscript; P.K. assisted in GEM study; N.C. and C.H. designed, performed LX47 PDX study and analyzed the data. DT, J.W., V.B., N.H., Y.Y. and U.D. performed some of the experiments; P.Z. and X.Z. synthesized and purified DT2216 and prepared the vehicle and formulated DT2216 for the studies; W.Z., shared resources, revised and commented on the manuscript; L.J., provided some of the SCLC cell lines, revised and commented on the manuscript; G.Z. supervised the synthesis, purification, and formulation of DT2216, and revised the manuscript; R.H. revised and offered constructive criticism to help improve the manuscript; M.Z.-K. and F.J.K. provided some of the reagents including SCLC cell lines and GEM model, revised and commented on the manuscript; D.Z. co-conceived and co-supervised the study, interpreted some of the data, provided resources and revised the manuscript. All authors discussed the results and commented on the manuscript.

