## Supplementary Information for "Co-targeting BCL-X_L_ and MCL-1 with DT2216 and AZD8055 synergistically inhibits small-cell lung cancer growth without causing on-target toxicities in mice"

### SUPPLEMENTARY METHODS

#### Immunoblotting

Cell pellets were lysed using RIPA lysis buffer (Cat. No. BP-115DG, Boston Bio Products, Ashland, MA) supplemented with protease and phosphatase inhibitor cocktail (Cat. No. PPC1010, Sigma-Aldrich, St. Louis, MO) as described previously[1]. Briefly, an equal amount of protein samples (20-40 µg/lane) were loaded to precast gels and transferred onto PVDF membranes. The membranes were blocked with 5% (w/v) non-fat dry milk in TBST buffer, and subsequently probed with primary antibodies overnight at 4 °C. After washing with TBST, the membranes were incubated with horseradish peroxidase (HRP)-linked secondary antibody for 1-2 h at room temperature. Finally, the membranes were incubated with chemiluminescent HRP substrate (Cat. No. WBKLS0500, MilliporeSigma, Billerica, MA), and were recorded using the ChemiDoc MP Imaging System (Bio-Rad, Hercules, CA). The immunoblots were quantified using Image J software. The primary antibody details are provided in Supplementary Table-1.

#### Colony formation assay

A total of 2,000 cells per well were seeded in 6-well plates. After overnight incubation, the cells were treated with AZD8055, DT2216, or a combination of the two for 10-14 days. Fresh treatment-containing medium was added to the plates every four days. At the end, the cells were fixed with methanol, and then stained with 0.1% crystal violet solution.

**Annexin-V/PI apoptosis assay using flow cytometry**

Cells were seeded in 12-well plates at a density of $1\times10^5$ cells per well and were treated with AZD8055, DT2216, or the combination of the two for 72 h. After incubation, the cells were collected by trypsinization and were stained with Annexin V-Alexa Fluor 647 (1:50, Cat. No. 640912, BioLegend, San Diego, CA) and propidium iodide (PI, 1 μg/mL, Cat. No. 421301, BioLegend) for 30 min at room temperature. The samples were analyzed using flow cytometer (Aurora, Cytek Biosciences, Fremont, CA) and the SpectraFlo software (Repligen, Waltham, MA). A minimum of 10,000 events were recorded for each sample. The percentage of apoptotic cells are presented as the sum of Annexin V-positive/ PI-negative and Annexin V-positive/ PI-positive cells.

**Histopathology**

Mouse lungs were fixed in 4% paraformaldehyde overnight and were then transferred to 70% ethanol. Paraffin embedding, sectioning and H & E stating were performed by the Molecular Pathology Core (University of Florida). The images were taken using snapshots from Leica Aperio Scanscope CS whole slide scanner (Leica Biosystems, Inc., Buffalo Grove, IL). The tumor area in each lung was calculated using QuPath Quantitative Pathology and Bioimage Analysis software (version 0.2.3) with Image J extension (https://qupath.github.io/).

### SUPPLEMENTARY FIGURES

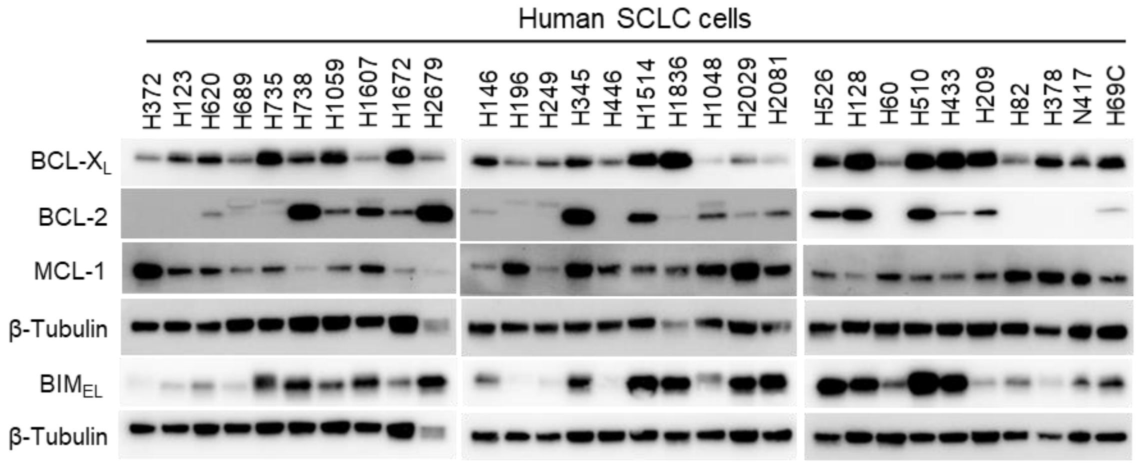


**Supplementary Fig. 1. The BCL-2 family of anti-apoptotic proteins are highly expressed in SCLC cells.** Immunoblot analyses of indicated BCL-2 family of antiapoptotic (BCL-$X_L$, BCL-2 and MCL-1) and proapoptotic (BIM-extra-large [$BIM_{EL}$]) proteins in 30 different human SCLC cell lines. β-tubulin was used as an equal loading control.

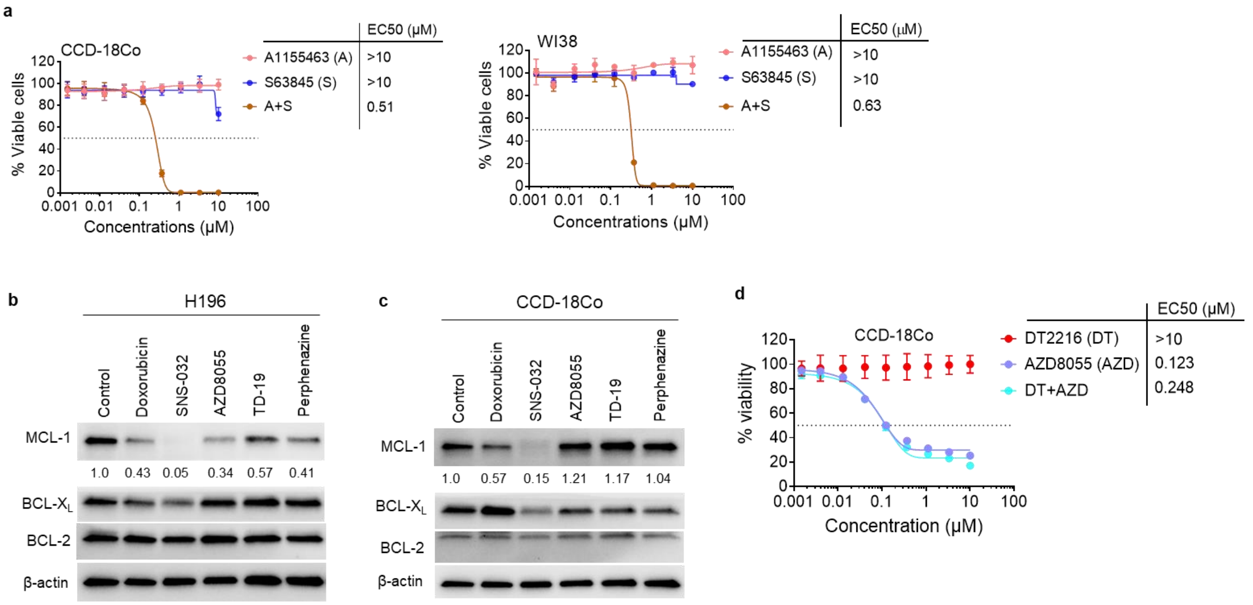


**Supplementary Fig. 2. AZD8055 selectively suppresses MCL-1 expression in tumor cells and its combination with DT2216 has no additive or synergistic activity in normal cells. a,** Viability of normal colon CCD-18Co and lung fibroblasts WI-38 cells after they were treated with increasing concentrations of DT2216 (DT) or S63845 (S) or their combination (1:1 ratio) for 72 h. **b, c**, Immunoblot analyses of MCL-1, BCL-$X_L$ and BCL-2 in H196 SCLC (b) and CCD-18Co cells (c) after they were treated with indicated agents (1 μM each) for 24 h. The normalized densitometric values for MCL-1 are shown underneath. **d**, Viability of normal colon CCD-18Co cells after they were treated with increasing concentrations of DT2216 (DT) or AZD8055 (AZD) or their combination (1:1 ratio) for 72 h.

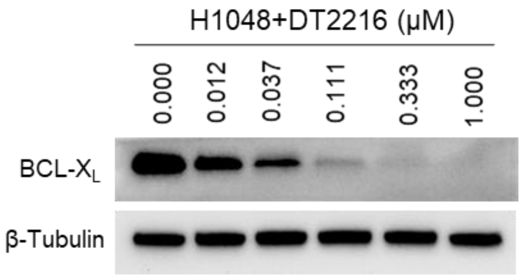


**Supplementary Fig. 3. DT2216 causes dose-dependent degradation of BCL-X$_L$ in SCLC cells.** Immunoblot analyses of BCL-X$_L$ in H1048 cells after they were treated with increasing concentrations of DT2216 for 16 h. β-tubulin was used as an equal loading control.

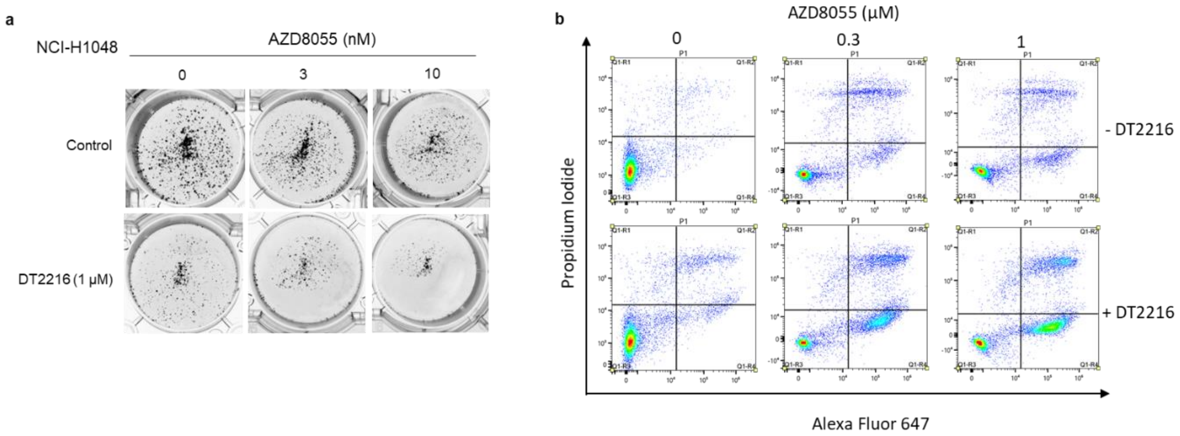


**Supplementary Fig. 4. AZD8055+DT2216 combination synergistically kills H1048 SCLC cells. a,** Images of colonies in H1048 cells after they were treated with indicated concentrations of DT2216 or AZD8055 or their combinations. **b,** Apoptosis analysis in H1048 cells after they were treated with indicated concentrations of AZD8055 alone or with DT2216 (1 µM) for 72 h.

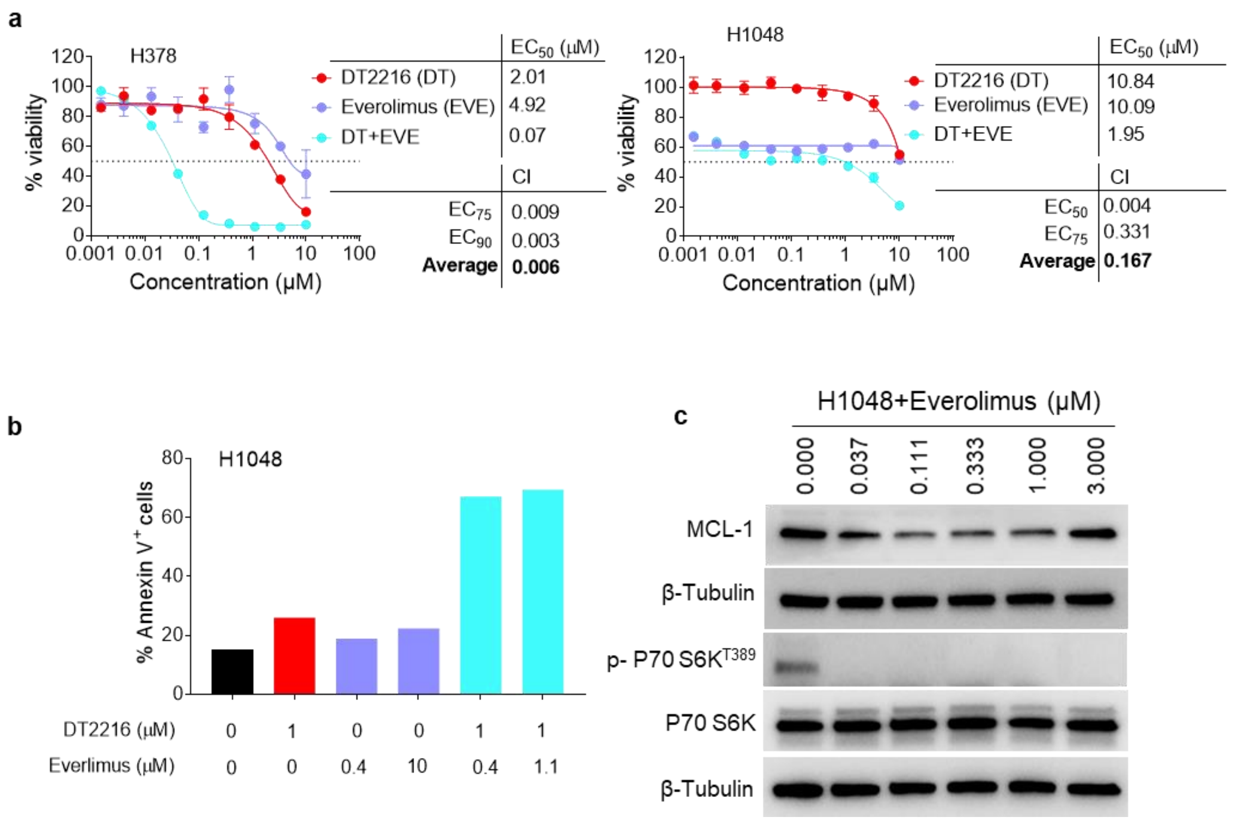


**Supplementary Fig. 5. Everolimus shows similar synergistic activity as AZD8055 when combined with DT2216 against SCLC cells. a,** Viability of H378 and H1048 SCLC after they were treated with increasing concentrations of DT2216 (DT) or everolimus (EVE) or their combination (1:1 ratio) for 6 days. $EC_{50}$ and CI values are shown. The data presented are mean ± s.d (n = 3 replicate cell cultures). **b,** Percentage of Annexin V+ (early + late apoptotic) cells after they were treated with indicated concentrations of DT2216 or everolimus or combinations for 48 h. **c,** Immunoblot analysis of MCL-1 and mTOR1 substrate P70 S6K after 24 h treatment with indicated concentrations of everolimus in H1048 SCLC cells.

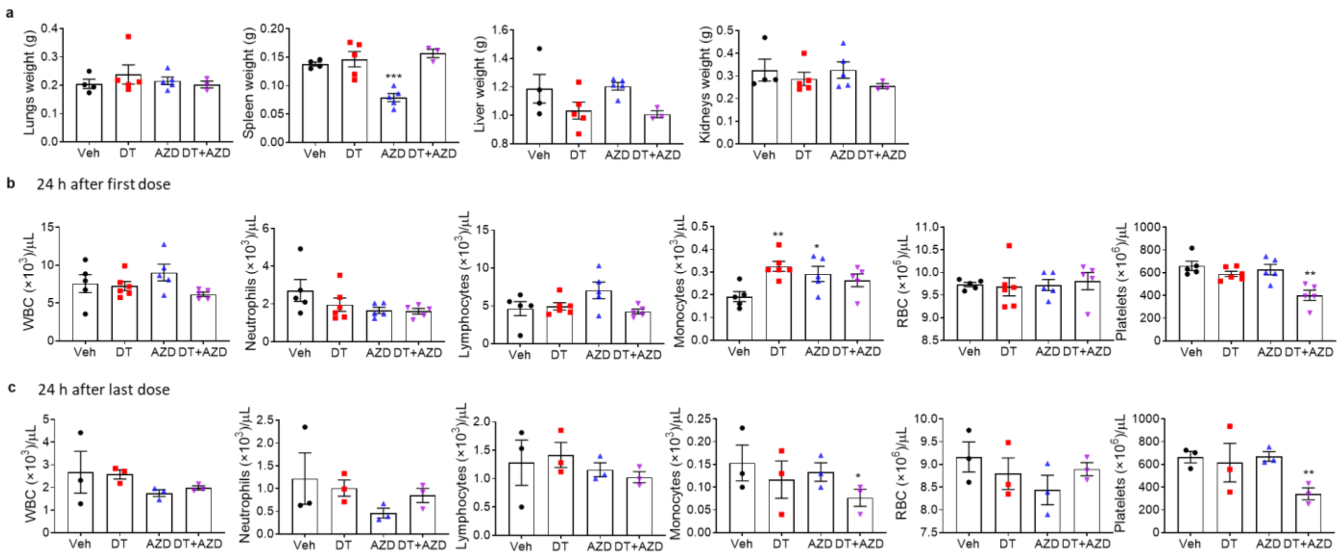

**Supplementary Fig. 6. The combination of DT2216 and AZD8055 causes no clinically-significant normal tissue or hematological toxicities in mice. a,** Weights of different organs harvested 24 h treatment with DT2216 (DT), AZD8055 (AZD) or their combination in *Rb1/p53/p130* mice as in **Fig. 6 e**. **b, c,** Counts of different blood cells 24 h after first dose (b) and 24 h after last dose (c) as in **Fig. 6 e**.

**Supplementary Table 1. Antibodies used in immunoblotting**

| Antibody | Clone | Antibody isotype | Catalog # | Concentration |
|---|---|---|---|---|
| BCL-$X_L$ | _ | Rabbit IgG polyclonal | 2762 | 1:1000 |
| BCL-2 | 50E3 | Rabbit IgG monoclonal | 2870 | 1:500 |
| MCL-1 | D35A5 | Rabbit IgG monoclonal | 5453 | 1:1000 |
| BIM | C34C5 | Rabbit IgG monoclonal | 2933 | 1:1000 |
| p-4EBP1 T37/46 | 236B4 | Rabbit IgG monoclonal | 2855 | 1:1000 |
| 4EBP1 | 53H11 | Rabbit IgG monoclonal | 9644 | 1:1000 |
| p-S6 S240/244 | _ | Rabbit IgG polyclonal | 2215 | 1:1000 |
| S6 | 5G10 | Rabbit IgG monoclonal | 2217 | 1:1000 |
| P-p70 S6K T389 | 108D2 | Rabbit IgG monoclonal | 9234 | 1:1000 |
| p70 S6K | _ | Rabbit IgG polyclonal | 9202 | 1:1000 |
| Full-length PARP | 46D11 | Rabbit IgG monoclonal | 9532 | 1:1000 |
| Cleaved PARP | D64E10 | Rabbit IgG monoclonal | 5625 | 1:1000 |
| full-length caspase-3 | _ | Rabbit IgG polyclonal | 9662 | 1:1000 |
| cleaved caspase-3 | _ | Rabbit IgG polyclonal | 9661 | 1:1000 |
| β-tubulin | _ | Rabbit IgG polyclonal | 2146 | 1:3000 |
| β-actin | D6A8 | Rabbit IgG monoclonal | 8457 | 1:5000 |
| Secondary antibody | | Anti-rabbit IgG, HRP | 7074 | 1:3000 |

**Footnotes:** All the antibodies were purhcased from Cell Signaling Technology, Danvers, MA.

### SUPPLEMNTARY REFERENCES


1. Khan S, Zhang X, Lv D, Zhang Q, He Y, Zhang P, et al. A selective BCL-XL PROTAC degrader achieves safe and potent antitumor activity. Nat Med. 2019;25(12):1938-47.
